# Constructing a consensus serum metabolome

**DOI:** 10.1101/2025.05.07.652782

**Authors:** Yuanye Chi, Joshua M. Mitchell, Maheshwor Thapa, Shujian Zheng, Zackary Frohock, Yi Li, Aleksandr Smirnov, Xiuxia Du, Shuzhao Li

## Abstract

Blood analysis is the most common in biomedical applications and a reference metabolome will be critical for effective annotation and for guiding scientific investigations. However, compiling such a reference is hindered by many technical challenges, despite the availability of large amount of metabolomics data today. Based on a new set of data structures and tools, we have assembled a consensus serum metabolome (CSM) from over 100,000 mass spectrometry acquisitions of more than 200 million spectra. This provides a comprehensive survey of human blood chemistry, revealing the frequency dependent nature of metabolome and exposome.

Major gaps are found between CSM and the current databases. The CSM enables community-level data alignment and significantly improves annotation quality of LC-MS metabolomics.

**Highlights:**

- A reference of human biochemistry linked to observation frequency
- Major gaps revealed in current databases and experimental methods
- Enabling cross-laboratory, cross-platform data alignment
- Accelerated and cumulative metabolite annotation

## Introduction

Human biochemistry is mostly operated via small molecules and metabolomics is the technology that aims to measure them comprehensively. Because the small molecules are chemically diverse, measurements are often method dependent. Untargeted methods strive to achieve high coverage of such measurements, while cross-laboratory data comparison is still difficult. There have been longstanding questions on how big the human metabolome is, how much untargeted experiments measure and how many metabolites are unknown. A reference metabolome, like the reference human genome, is needed to align metabolomics data from around the world, towards effective annotation and guiding scientific investigations. However, compiling such a reference is hindered by many technical challenges, despite the availability of large amount of metabolomics data today. Since blood reflects the circulating metabolites in human bodies and blood analysis is the most common in biomedical applications, a reference serum metabolome is the most important first step.

The first serum metabolome database was reported in 2011 by Wishart and colleagues, who employed 5 platforms to measure 3564 compounds. Of those compounds, 3394 were from lipid profiling (Psychogios et al, 2011). This database was later merged into the Human Metabolome Database (HMDB, Wishart et al, 2022). As the technologies evolved, LC-HRMS (liquid chromatography coupled high-resolution mass spectrometry, LC-MS herein) has become dominant in metabolomics assays, often annotated by authentic chemical standards and MS/MS libraries. Bar et al (2020) described a reference serum metabolome, based on 1,251 metabolites measured by a commercial vendor using four LC-MS methods. Of these compounds, 411 were lipids, 498 likely xenobiotics (including 295 unknown compounds), 44 peptides, while the remaining 298 metabolites covered amino acid, nucleotide, carbohydrate and cofactor pathways. Metabolic pathways are often combined into genome scale metabolic models (GSMM). A representative human GSMM contains about 4,000 metabolites without considering their intracellular compartmentalization (Robinson et al, 2020; Brunk et al, 2018). In general, there is a great discrepancy between these databases and experimental data. About 15% measured intracellular metabolites are covered by GSMM (Mitchell et al, 2024b). About 70% of common metabolomics data are not found in the databases (Metz et al, 2024; Uppal et al, 2016; El Abiead et al, 2025; Chi et al, 2025). Therefore, a reference metabolome should be based on broad survey of community data.

Historically, the assembly of genomes built on genetic maps and sequence alignment algorithms; the annotations cumulated in databases over time, including gene structures (e.g. exons and introns) and functions. The assembly of metabolomes also requires a set of data models and algorithms, but they are very different from genomics, because metabolites have diverse chemical properties, and their mass spectrometry data are not ready to be digitalized into 4 nucleotides (genomics) or 21 amino acids (proteomics). In this study, we have designed a series of data structures and tools to enable a first draft of assembling a consensus serum metabolome (CSM) from large-scale public data. The results provide a new data driven survey of human blood chemistry. The CSM provides a framework for community level data alignment and cumulative annotation. We demonstrate the power of CSM by biochemical phenotyping of patients across cohorts.

## Results

### Consensus mass registries from large-scale analysis of LC-MS metabolomics data

LC-MS metabolomics data of human serum or plasma samples were collected from public repositories (**Figure 1A, Method**). This included over 100,000 acquisition files from 136 studies using the orbital mass spectrometers. In addition, 26,253 acquisition files from the TOF (time of flight) mass spectrometers are analyzed in order to compare to the orbital results. Each dataset contains thousands of features, which are defined by elution profiles in consecutive spectra. The key of assembling them into a useful reference resides on data structures to model different levels of data, including m/z (mass to charge ratio), separation of isomers, ion relationships and putative compounds (**Suppl Figure 1A**). Many mass spectrometers used in metabolomics today can achieve a mass resolution of 1∼5 part per million (ppm), which is adequate to resolve most chemical formulas of smaller molecules. The m/z values are therefore comparable cross labs, while the chromatography may differ greatly. The first step of assembling CSM is to define the common mass values.

**Figure 1:**
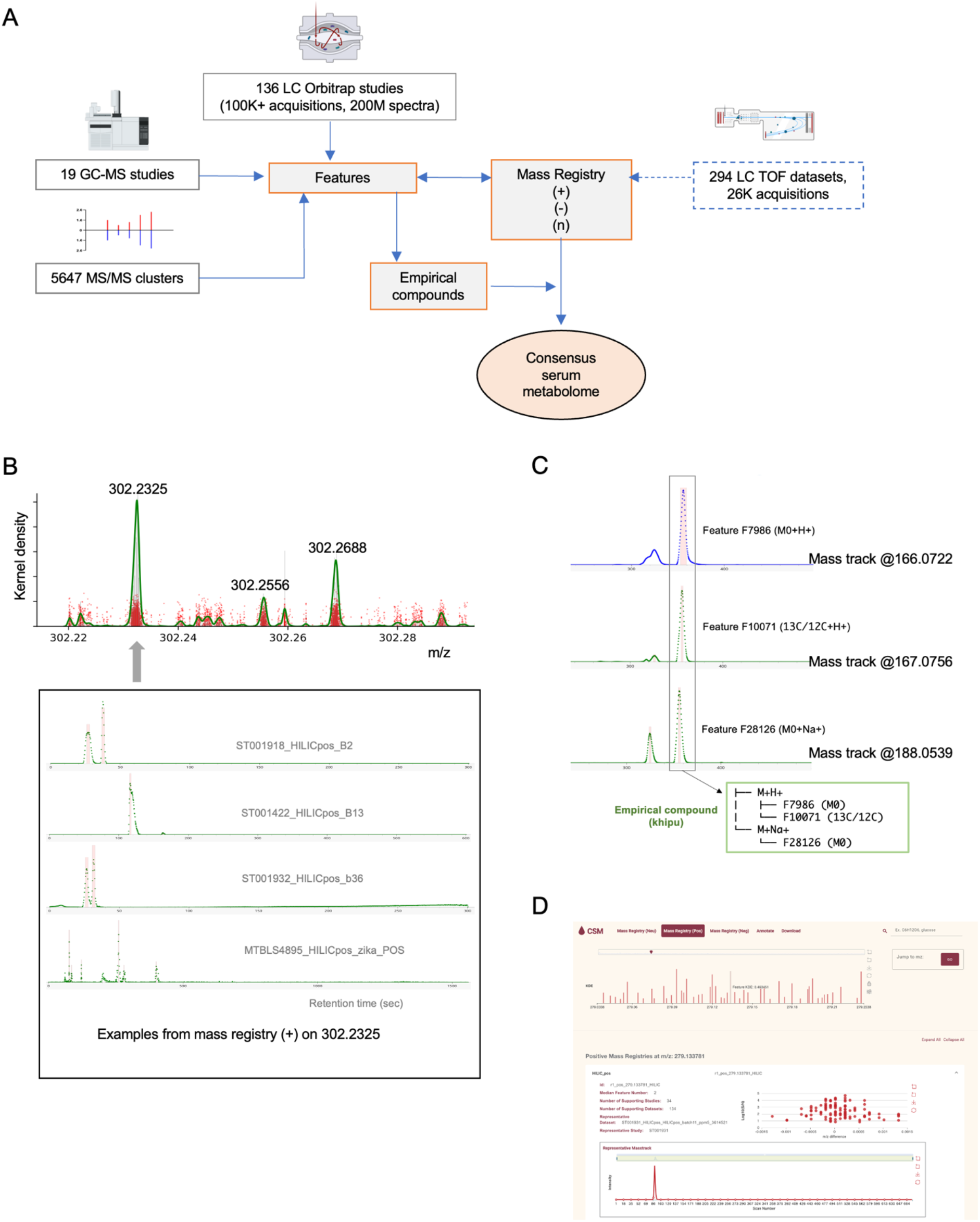
Assembly of consensus serum metabolome. A) Schematic of data sources and assembly process of CSM. LC-MS metabolomics data of human serum or plasma samples were collected from public repositories. They were processed into features defined by m/z and retention time. Most common m/z values were selected as mass registries. Pre-annotation organizes isotopologues and adducts into empirical compounds, within each dataset. GC-MS and MS/MS data were integrated with compound annotations. The CSM contains neutral mass registries and associated consensus features; each of the latter has consensus m/z value and retention time index. Details are provided in Suppl Figure 2. B) Consensus mass registries are based on KDE peaks from all datasets. Mass tracks are reported in each dataset and their m/z values are centered on distinct values. Each red dot in the first KDE plot is one feature reported by a dataset. Over 1000 entries contributed to peak 302.2325. The mass tracks have varying number and quality of elution peaks, which are used in majority vote for a consensus number of features in CSM. C) Illustration of pre-annotation, grouping ions into an empirical compound (khipu) within a dataset. D) Screen shots of CSM website (https://metabolomics.cloud/csm) for navigation and annotation.

For a common metabolite, m/z values around the theoretical value are reported by many datasets (**Figure 1B**, red dots in the upper panel). These collective statistics can be extracted by kernel density estimation (KDE), and the KDE peaks become mass registries (**Figure 1B, Suppl Figure 1B**). By a KDE threshold of 5% of maximum, 48,697 consensus mass registries are identified in the positive ionization mode, and 40,483 in the negative mode (**Suppl Figure 2**). Compared to the 1,251 metabolites reported by Bar et al (2020), 95% of them are matched in our mass registries. Supporting each mass registry are mass tracks from many datasets (**Figure 1B**). The number of elution peaks on a mass track depends on the number of isomeric compounds, experimental method and data quality. The consensus number of features per LC category per mass registry in CSM is determined by majority vote of all supporting datasets.

From the positive and negative mass registries, neutral mass values are calculated by pre-annotation using the khipu algorithm (Li and Zheng, 2023). Pre-annotation organizes isotopologues and adducts into empirical compounds (**Figure 1C**). This is performed within each dataset due to the requirement of coelution of these ions. Because a CSM feature is linked to voting features in individual studies, the over 6 million ion relationships from all datasets can be transferred into CSM to assign ion types and determine neutral mass values (**Suppl Figure 2**). The result shows that CSM features are mostly [M+H]+ or [M-H](36%) and 13C isotopes (25%). This is because the construction is frequency driven and other adducts are less frequent/consistent across studies. We have previously shown that in-source fragments are ∼3% of a typical LC-MS feature table (Chi et al, 2025). Their presence in CSM is expected to be rare. Together, this release of CSM consists of 69,677 neutral mass registries and 231,349 CSM features. They are accessible via an open web tool for search, navigation and annotation (**Figure 1D**).

### CSM as a frequency dependent overview of blood biochemistry

Human biochemistry is mostly based on six elements, carbon, hydrogen, nitrogen, oxygen, phosphorus, and sulfur (CHNOPS). The ratios of atoms in a molecule, e.g. hydrogen/carbon and oxygen/carbon, are useful to chemical analysis and visualization (Brockman et al, 2018). Fossil fuels often have O/C ratios under 0.1. The O/C range of CSM compounds is mostly between 0 and 1. The H/C ratios of the majority of CSM compounds fall between 0.5 and 2.5, those for lipids typically in 1.5 to 2.2. The O/C ratios of lipids are typically under 0.6, while nucleotides and sugars contain more oxygen. The full range of CSM compounds is visualized using a circular van Krevelen plot (Li, 2025), where the angular axis is H/C ratio and radial axis O/C ratio (**Figure 2A, Method**). Only about 1,000 compounds are detected in over 50% of the datasets (less at sample level), ∼4,000 over 20% and ∼10,000 over 10% datasets (**Figure 2A**, right). We separate the CSM compounds by their detection frequency: those present in < 10% datasets are plotted in the outer circle, and the more frequent compounds in the main middle circle. Thus conceptually, the outer circle represents exposome (Vermeulen et al, 2020; Rappaport et al, 2014) and the middle circle metabolome. This rough distinction will become better defined as more annotation is available.

**Figure 2:**
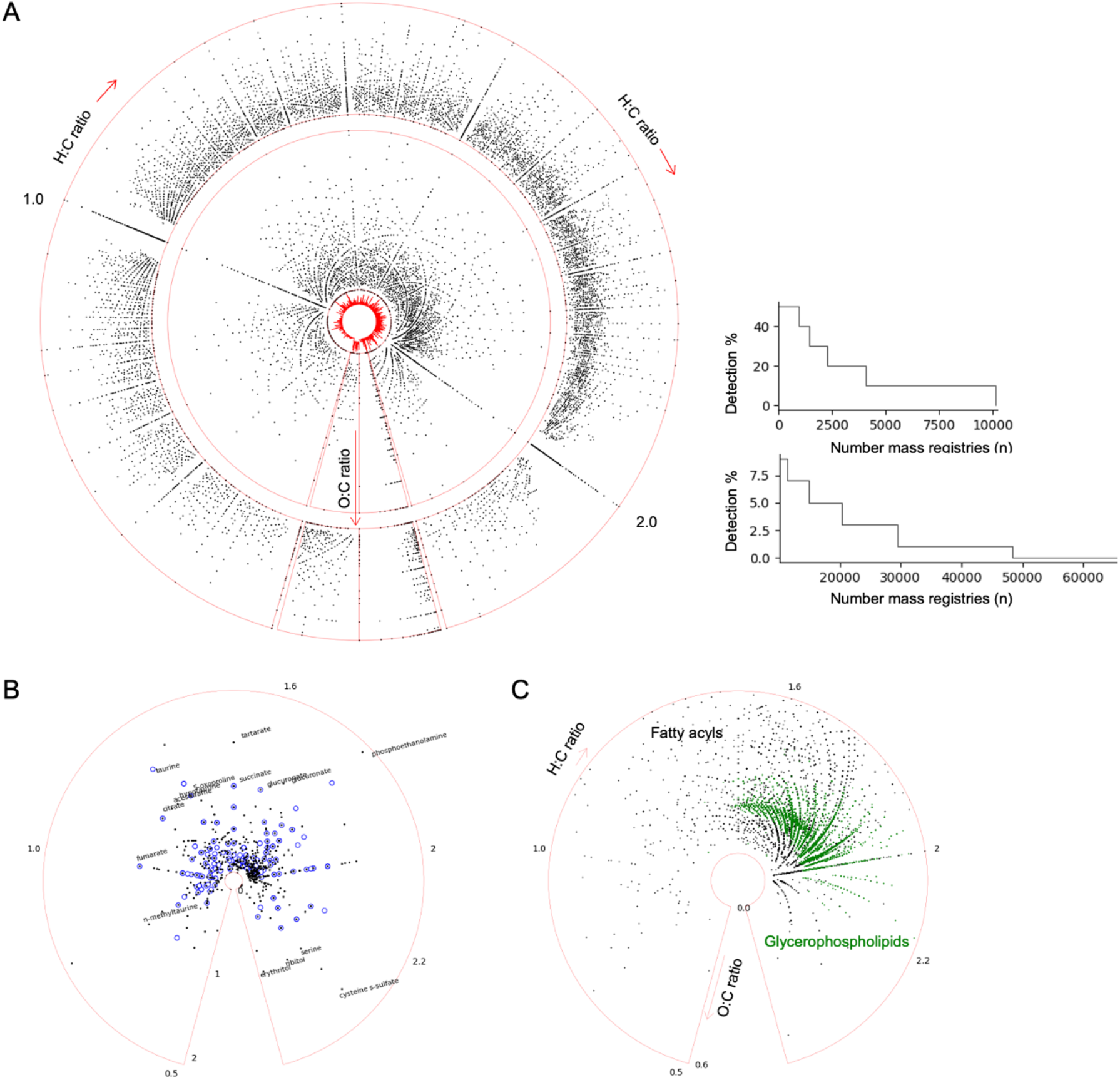
Overview of CSM composition. A) Each dot is a chemical formula in CSM. The polar axis is hydrogen/carbon ratio and radial axis oxygen/carbon ratio. The outer circle contains compounds detected in less than 10% in all datasets; middle circle for the more frequent compounds. The small center circle is a histogram of frequence of observations. Frequency distribution by percentage steps is shown on the right. B) Visualization of two authentic compound libraries on the same polar scales. C) Two lipid classes from HMDB are plotted to illustrate regular patterns.

The metabolomics community depends on authentic compound libraries for annotation. We were able to verify within CSM 589 compounds from the library underlying Bar et al (2020, commercial vendor) and 173 compounds from the library in Pathmasiri et al (2024, academic lab). These are representative numbers in the field, considering that some compound libraries are bigger but not specific to human blood. While neither comes close to be representative of the size of CSM, their projection on the same circular van Krevelen plot also shows their differences in selecting the library compounds (**Figure 2B**), which are reflected in coverage and method differences. Distinct curve patterns can be seen in **Figure 2A** because of organizational principles in chemical structures. For example, many of the curve patterns are driven by certain lipid classes (**Figure 2C**).

### Platform biases and major coverage gaps in current data landscape

The most common methods in LC-MS are positive or negative electrospray ionization (ESI) coupled with reverse phase (RP) or hydrophilic-interaction chromatography (HILIC). The use of chromatography has major impact on what compounds are detected (**Figure 3A**). When different studies are clustered by overlap compounds, two groups are separated by positive or negative ionizations (**Figure 3B**). Strikingly, the average overlap of compounds between two studies in **Figure 3B** is 8% (using Jaccard index, compound annotation explained in a later section). Using the features matched to CSM, the average overlap between two studies is 11%. These results indicate that the platform biases should inform future study design and data interpretation; most detected features are currently not reproducible cross laboratories and studies.

**Figure 3:**
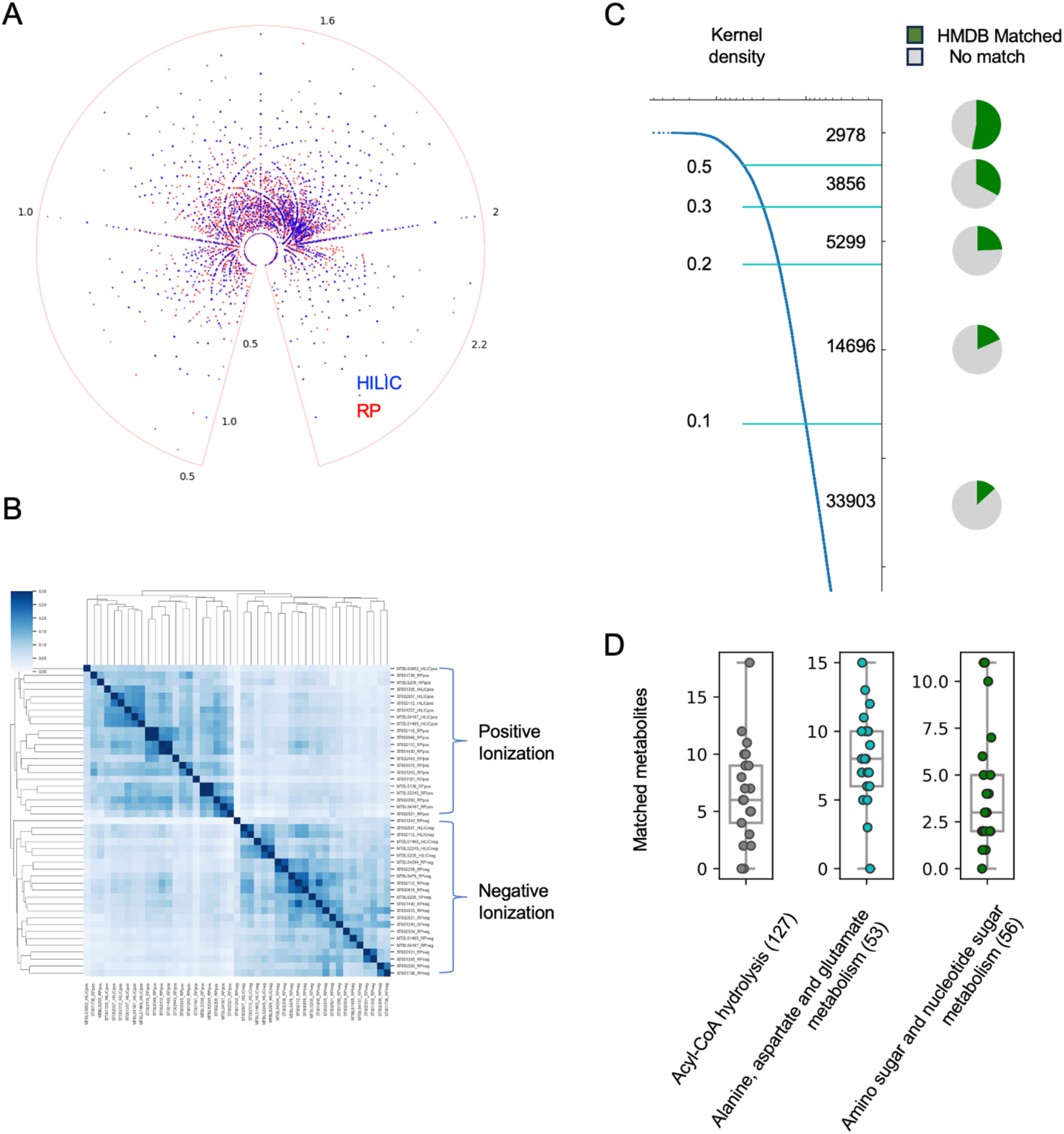
Platform biases cross datasets and gaps compared to current databases. A) Two major chromatography types are colored separately, on the same metabolome visualization in Fig. 2A. B) By overlap neutral mass values using Jaccard index, unsupervised hierarchical clustering separates positive and negative ionization datasets. C) Overlap between CSM and HMDB is dependent on compound detection frequency. KDE values and corresponding neutral mass numbers are marked on the left, and pie charts show overlap in each segment. D) Detection of metabolites on example pathways using a single HILIC ESI+ method. Each dot is from one dataset.

The CSM is compiled from many public datasets, while the existing metabolite databases are mainly based on literature or chemical standards. We can now compare CSM to existing databases via neutral mass, which is roughly at the level of chemical formulas, even without confirmed metabolite identity. While this approach does not completely confirm a positive match, it confirms a negative match. That is, if its neutral mass is missing in a database, the compound itself is missing in a database. In the version 5 of HMDB, 220,945 molecules are cataloged, by literature records and spectral data from chemical standards. The comparison between CSM and HMDB reveals that 53% of the 2,978 most frequent detections in CSM (KDE > 0.5) are matched to HMDB, while the overlap decreases greatly for the less frequent detections. Overall, about 80% of CSM is not found in HMDB (**Figure 3C**).

A representative collection of human metabolic pathways (Robinson et al, 2020) contains 3384 metabolites, which covers about 14% of the 2,978 most frequent detections in CSM (KDE > 0.5). This overlap decreases greatly for the less frequent detections (**Suppl Figure 3A**).

Conversely, the CSM covers about 60% of the metabolites in the pathway collection (**Suppl Figure 3B**). However, some pathways are poorly covered, and a single method often covers a small fraction of a metabolic pathway (**Suppl Figure 4, Figure 3D**). Strong method bias exists, e.g., carnitines are mostly detected in positive ionization (**Suppl Figure 4**). Even though blood is not the primary tissue for studying metabolic pathways, the coverage biases should be considered in the interpretation of metabolomics data.

### CSM enables data alignment cross studies

Cross-laboratory data comparison is a longstanding roadblock for metabolomics. Numerous ring trials on common materials have been conducted. Even though reproducibility can be excellent on standardized methods, and data alignment is feasible therein (Bonini et al, 2020, Habra et al, 2024), new method development and a common reference are required to align data from broader communities. Because a) metabolite identification is far from definitive (Metz et al, 2024), and b) the current metabolomic atlas projects do not handle unknown compounds (Rakusanova et al, 2023), the CSM is the first practical solution.

As already suggested in **Figure 3B**, reproducibility is limited cross studies. A top reason is that each dataset contains only a small subset of high-intensity, high-quality features (**Figure 4A**). This subset can still differ between laboratories, given method variation and optimization for different metabolites of interest. We have previously addressed the variations in computational software processing comprehensively (Li et al, 2023). The analysis here focuses on the cross-study alignment problem and analytical variations, starting with the same material analyzed by the same lab (**Figure 4B-C**), then onto the same material analyzed by different labs (**Figure 4D**), and different large cohorts analyzed by different labs (**Figure 4E**).

**Figure 4:**
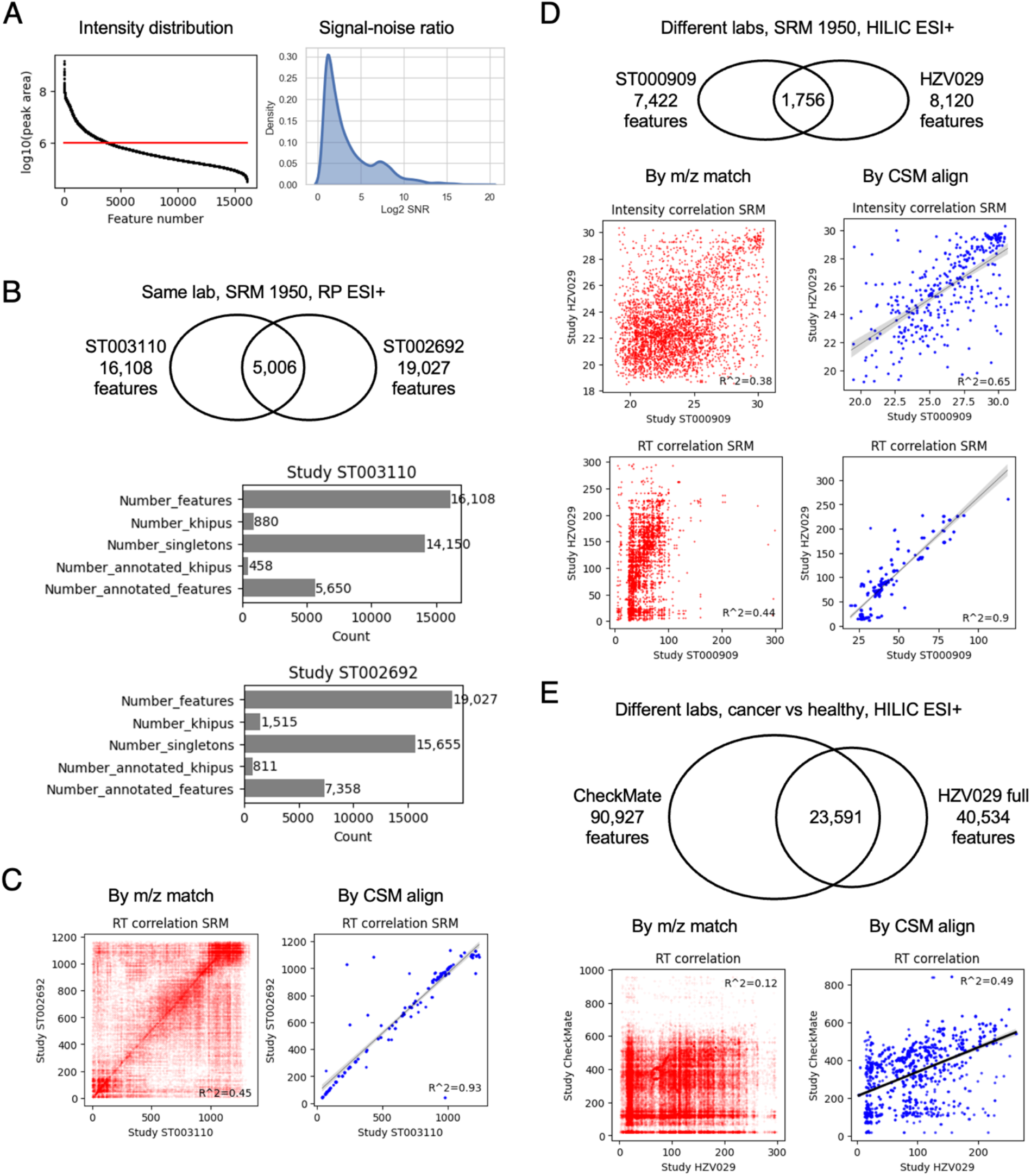
Alignment of metabolomics data cross labs via CSM. A) High intensity (left) and high quality (right) features are minority in each dataset. B) The NIST reference material SRM 1950 analyzed in two metabolomic experiments by the same lab. The SRM 1950 samples were part of studies ST003110 or ST002692 on Metabolomics Workbench. Top: overlap m/z values. Bottom: pre-annotation and CSM annotation of data. Khipus are putative compounds with patterns of multiple ions. Singletons are features with no relationship found to other features. Annotated khipus are the high-confidence portion of the data. C) Alignment of the above two datasets. Results in red are from features matched by m/z only; blue by CMS alignment. Because retention time (RT) is metabolite specific, RT correlation indicates success of alignment. D) The same reference samples analyzed by different labs. E) Alignment of a cancer study from Broad Institute (CheckMate, 1377 samples. Li et al, 2019) and our full HZV029 study (vaccination cohort, 1685 samples). Note that different chromatography between labs dictates the possible level of RT agreement.

The SRM 1950 is a pooled human serum sample, distributed by the National Institute of Standards and Technology (NIST), commonly used as a reference material by many labs (Phinney et al, 2013). The datasets ST003110 (Pathmasiri et al, 2024) and ST002692 (Taibl et al, 2023) were analyzed by the same lab in the NIH HHEAR network, using the same method (ESI+, PR), and included the SRM 1950 sample in several replicates. When the SRM 1950 replicates are compared between these two datasets, less than 1/3 mass values are in common (**Figure 4B**). For each dataset, the features are grouped into khipus (empirical compounds, group of ions belonging to same compound) in the pre-annotation process. E.g. 16,108 features in ST003110 are organized into 880 khipus and 14,150 singletons. The singletons are less reliable because they have no observed isotopologues. The number of khipus reflects the number of reliable compounds (**Figure 4B**). Even when m/z values are matched, many features are not the same metabolites, indicated by the plot of retention time, since RT is reproducible in the same lab (**Figure 4C**, left). Using the CSM alignment, the identical metabolites are extracted from the two datasets, as indicated by their consistent RTs (**Figure 4C**, right) and intensities (**Suppl Figure 5A**). The alignment of two other studies by a different lab, using HILIC chromatography, is shown in **Suppl Figure 5B**. RT correlation here is an independent test of CSM, because CSM alignment does not use study specific RT directly. The CSM alignment benefits from the empirical statistics in the reference metabolome, and from the understanding of feature relationships in the whole dataset (**Methods**).

We next compare the SRM 1950 replicates between two different labs, ST000909 study by a lab in the NIH CHEAR network and HZV029 study from our lab. The successful alignment of common metabolites is demonstrated by their consistent intensities and RTs (**Figure 4D**). We note here that the chromatography is quite different between the two labs, but a linear relationship is recovered by CSM alignment.

Data of reference materials are not always representative of real biomedical studies, especially of disease cohorts. We next apply CSM to align two large datasets, a cancer study from Broad Institute (CheckMate, 1377 samples. Li et al, 2019) and our full HZV029 study (vaccination cohort, 1685 samples). The two studies share a high number of m/z matches (**Figure 4E**), because their large sample sizes make many infrequent compounds detectable. Again, m/z match does not warrant correct match of metabolites. A clear positive correlation of RT emerges after CSM based alignment, suggesting that identical metabolites are found. Important difference of chromatography prevents precise RT matching (12 minutes for CheckMate, 5 minutes for HZV029), but the correlation is validated using limited number of common metabolites that were identified using authentic chemical standards in both labs (**Suppl Figure 6A**). Taken together, the above results indicate that CSM allows interlaboratory data alignment from similar methods. CSM features are linked to neutral mass registry and empirical compounds, which are generic to any platform and methods. When new data and new annotations are available, they can be easily integrated into the reference.

### A reference platform for reusable and cumulative annotation

An important function of CSM is to annotate metabolomics data, at the levels of both pre-annotation and compound identity. Pre-annotation computes the relationships between features (**Figure 1C**). After a feature is determined to be part of a khipu, it will not be mistaken as a new compound. Thus, the number of confidently measured compounds in each experiment can be calculated (**Figure 4B**). A systematic analysis of all public LC-MS studies used in CSM shows that 30,000∼40,000 features are measured in an experiment, while khipus of good isotopic patterns range from 840 for HILIC ESI-experiments to 1663 for RP ESI-experiments (**Suppl Figure 7A**). The numbers of matched mass tracks indicate that the majority of mass tracks contain only one elution peak, an important factor in compound identification.

Since the CSM features are voted by features in individual studies, both the pre-annotation and annotation of a voting feature can be carried over to CSM. In a typical CSM feature, the type of ion is determined by majority vote (**Suppl Figure 7B**). A direct benefit is that a high number of singletons in each study gains pre-annotation by matching to CSM features, e.g. the unknown ions can be assumed as [M+H]+ in **Suppl Figure 7B**. They also gain detection confidence because the feature has been observed in over 200 other datasets. Most studies have over 10,000 features that match to CSM (**Figure 5A**), and over 2,000 as confident compounds (khipus, **Figure 5B, Suppl Figure 6B**).

**Figure 5:**
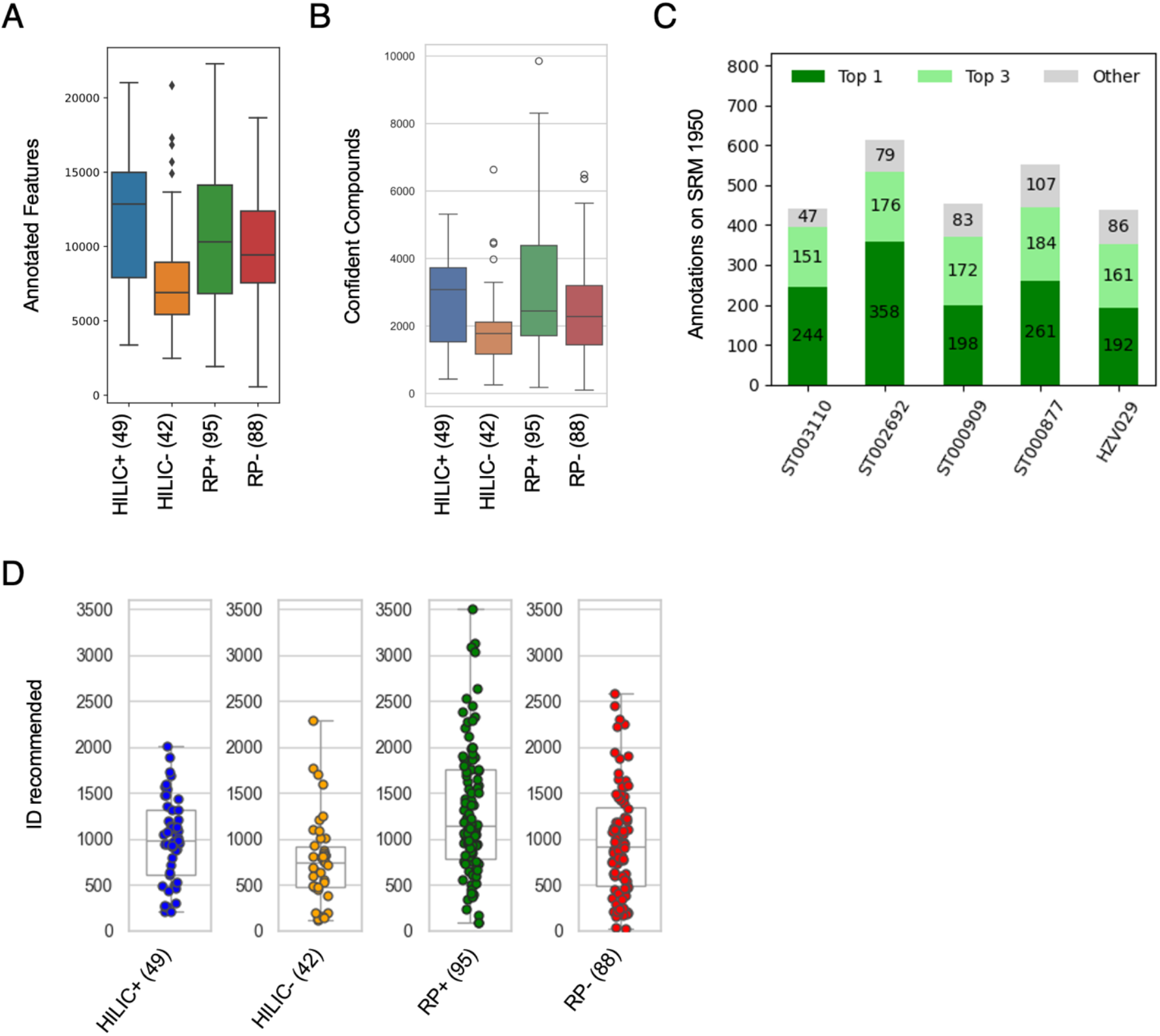
Annotation coverage and quality on public datasets by CSM. A-B) Numbers of annotated features and high-confidence compounds aligned to CSM in 244 datasets of four methods, regardless of identification. C) CSM compound annotation on 5 datasets of NIST SRM 1950, compared to the report by Mandal et al, 2025. Dark green is number of agreements in top CSM recommendation; light green agreement in top 3 CSM recommendation. D) Numbers of named annotated compounds on the same 244 datasets as in A).

Authentic compound libraries are the gold standard in metabolomics annotation, but they are platform specific and few of them are released in the public. Reference retention time is introduced for a larger number of compounds from the NORMAN (RP chromatography, Aalizadeh et al, 2021) and RETIP (HILIC chromatography, Bonini et al, 2020) databases. A regression procedure, similar to the RT alignment in Asari (Li et al, 2023), is used to remap the RT in each study to CSM; in return, a consensus RTI (retention time index) is taken from the median value of all contributing features per CSM feature. This RTI is not precise or specific enough to replace a lab specific library, but useful to resolve elution orders in CSM annotation.

A blood compound database was compiled from the Blood Exposome Database (Barupal and Fiehn, 2019) and the serum subset of HMDB, and augmented by verified compounds during the CSM construction. When new data are compared to CSM, isomers may need to be resolved and a CSM feature can match to multiple database entries. A set of heuristic algorithms were implemented to prioritize recommendations by prior identification, detection frequency, partition coefficient, reported compound concentration in human blood, and prior publications. The compound annotation by CSM returns a top recommendation and scores of all probable identities. Hence, annotation of a dataset by CSM is based on a reference metabolome, taking into account the relationships between features in a defined biological tissue.

A set of annotation of NIST SRM 1950 by multiple analytical platforms was released recently by Wishart and colleagues (Mandal et al, 2025), therefore can serve as a benchmark of the annotation quality by CSM. The five SRM 1950 datasets in **Figure 4** and **Suppl Figure 5** are annotated by CSM to receive recommended identities per features. The results show high degrees of agreement with the annotation provided by Mandal et al, 2025 (**Figure 5C**), i.e., assign high-quality annotation of 300∼400 compounds per dataset per method. The total feature numbers are small in these small SRM 1950 datasets. The CSM annotates higher number of compounds in larger studies (**Figure 5D**). E.g., of the CheckMate data, 5,286 compounds have high-confidence CSM matches, of which 1,872 have recommended compound identities, significantly more than 202 annotated metabolites in the original paper. Compared to the authentic compound library provided by the original analytical lab, good agreement is observed for the overlap compounds (**Suppl Figure 6C**). These CSM annotated compounds are meaningful, as they can accurately predict patient sex and treatments using machine learning tools (**Suppl Figure 6D**).

Building authentic compound libraries happens in every metabolomics lab therefor a lot of repetition worldwide. These results from CSM show that a reference metabolome can share annotation in high degree of confidence, even for LC-MS data. Using the software package or web tool of CSM, high confidence features and associated annotations can be obtained for a dataset in minutes. This is not dependent on but can be complement to a lab specific compound library.

### Computational standardization of biochemical phenotyping

The CSM and algorithms are designed for consistence within a dataset and cross the reference metabolome. This builds standardization into the CSM annotation results. As such, this removes many complex issues in searching mass spectrometry data and comparing data cross studies.

For example, CSM feature r1_neg_498.930239 is perfluorooctane sulfonic acid (PFOS). A string search of r1_neg_498.930239 finds that a number of samples in study ST001932 have detectable levels of PFOS. Since consistent CSM identifiers are used, cross-study analysis is now feasible, even if compounds are unidentified or have inconsistent names. To illustrate such applications, we examine four common compounds in five studies (**Figure 6A**). The detection of caffeine is very common. Cotinine, a metabolite of nicotine, is detected at trace level in many samples, suggesting that the few samples of high intensity are true positives. The study ST001932 is a pediatric diabetic cohort, explaining that a) the caffeine level is lower than adult cohorts, and b) the diabetic drug metformin is detected in a number of samples. Salicylic acid is a metabolite of aspirin, detected in the two studies that were part of a cohort on aspirin treatment (Barry et al, 2022).

**Figure 6:**
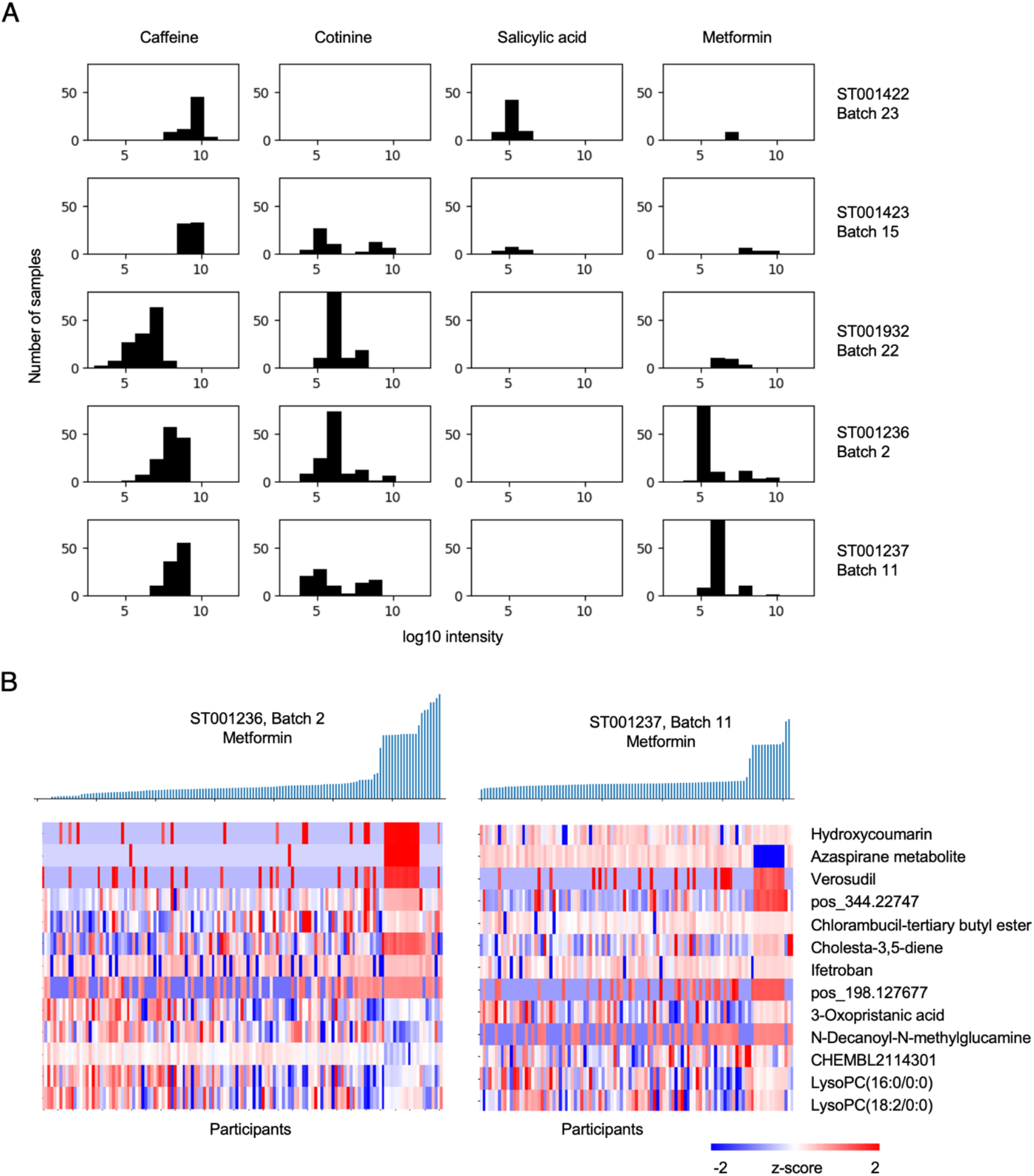
Biochemical phenotyping on CSM standardized datasets. A) Detection of four compounds in five datasets. Intensity (x axis) values of 1E5 is considered low on these platforms, which do not have the same standards. B) Distribution of metformin intensity in two studies (top) and their correlated metabolomic features (bottom). Each column is one patient.

Metformin is also detected in ST001236 and ST001237 (**Figure 6A**), two independent renal cell carcinoma cohorts (Li et al, 2019). The low detection level in most samples suggests basal contamination in the analysis, which can be controlled by blank samples in the experiments if provided. Nonetheless, since high metformin levels exist in some samples, we can ask what metabolomic features are correlated with metformin consistently in these two cohorts (**Figure 6B**). The features most correlated with metformin here appear to be prescription drugs. All patients had documented cancer treatments, but this result indicates that a subgroup of patients required medications for other complications. This type of information will be valuable for understanding patient responses, especially when clinical record is not available on legacy data. Inversely correlated to metformin include two lysophosphatidylcholines, corroborating a known therapeutic effect of metformin lowering lysophosphatidylcholines (Wanninger et al, 2008; Cai et al, 2009). These results indicate that CSM provides a standardized platform that can support speedy analysis and cross-study interpretation.

## Discussion

The CSM is a reference metabolome that is representative of community-wide data. The results reveal that a great number of compounds are missing in the current metabolite and metabolic databases; and conversely, many reported metabolites are not measured by current metabolomic experiments, especially on a single method. The large number of low-frequency compounds suggests the ubiquitous presence of exposome in human data, and challenges the gene-centric view of biochemistry.

The CSM is different from traditional metabolite databases, because it archives a large number of unknown compounds; the detection frequency is tissue specific; and it supports alignment and annotation of full datasets. The latter utilizes the distribution and relationships between features, considering platform bias. Using genomic terms, CSM supports “global alignment” while only “local alignment” is available in metabolite databases. The CSM differs from atlas projects, by handling the unknown and by including diverse platforms and data that cannot be covered by a few labs.

The benefits are immediate. Untargeted metabolomics produce features of varying quality and consistency. A metabolite that is missed or measured in low quality in one study may be measured in high quality in another study. By leveraging a consensus reference, confidence can be assigned to features more accurately. This is evident in the several thousand features supported by CSM in a typical human cohort study (**Figure 5**). The computational annotation of core metabolites was validated on several SRM 1950 datasets. This means any laboratory facility in the world can obtain state-of-art annotation on serum or plasma metabolomics, even without a big compound library, which is laborious and costly to build. More importantly, CSM is a framework that enables cross-laboratory, cross-study data alignment and cumulative annotation at the community level. Retrospective annotation is now feasible, which is important, given that most datasets in public repositories have no verifiable annotation.

Many improvements can be made on CSM. The current release only focused on major LC-MS platforms and did not include lipidomics-only experiments. Meta data are often not complete on the public datasets; controls are underutilized, including blank sample controls. However, comparison to known contaminants indicates few of them are in CSM. The CSM infrastructure allows new annotations to be added and shared. The quality and coverage of CSM annotation can grow significantly, given that initial annotations were only confirmed on two compound libraries. Majority vote is not always the correct result. Future efforts will include manual curation. Similar to the release cycles of the human genome assembly, the CSM can be continuously improved by new data and annotation packaged into new releases.

## Data Availability

The data release, notebooks for data analysis, and supporting software code are freely available at https://github.com/shuzhao-li-lab/consensus_serum_metabolome.

## Author Contributions

Project design and writing: SL. Data collection and processing: YC. In-house spectra acquisition: MT and SZ. Data analysis and software development: SL, YC and JM. Web tool development: YC. Collection and analysis of public GC-MS and MS/MS data: AS and XD. Machine learning of CheckMate data: ZF and YL. All authors participated in and approved the final manuscript.

## Acknowledgements

This work was in part supported by NIH grants U01 CA235493 (NCI), R01 AI149746 (NIAID), and ARPA-H award D24AC00345.

## Supplementary Information

Supplementary Figure 1: Data models and assembly of consensus mass registries. Supplementary Figure 2: Detailed construction process of CSM.

Supplementary Figure 3: Overlap between CSM and a genome scale metabolic model. Supplementary Figure 4. CSM coverage of pathways in human genome scale metabolic model by methods.

Supplementary Figure 5: Alignment of studies from the same labs. Supplementary Figure 6: Reprocess, annotation and analysis of CheckMate data. Supplementary Figure 7: Details of pre-annotation.

## Methods

### Data retrieval and processing

All metabolomic studies from the repositories, Metabolomics Workbench and MetaboLights, were retrieved by the following criteria: human serum or plasma samples; more than 50 samples; using LC coupled with Orbitrap or TOF based technologies; samples were not prepared solely for lipidomics; no data processing errors. The resulting data consist of 1086 datasets from 136 studies from Orbitrap platforms. A dataset is based on a single method. The majority datasets fall into either HILIC or RP chromatography, using either positive or negative ESI (electrospray ionization). No other column type was included in this version. Large studies were split into multiple datasets to reduce bias in processing and annotation. The splits were based on experimental batches, sized between 100∼120 samples. Because feature statistics at the dataset level are dependent on the number of samples. Processing software may have more errors in aligning very large datasets. By limiting dataset size, we avoid such potential issues.

The raw files were converted to centroided mzML files using ThermoRawFileParser v1.3.1 for Orbitrap data or msConvert v3 for TOF data. Data preprocessing was performed using Asari (v 1.13.1), yielding feature tables with m/z, retention time, peak shape, signal-to-noise ratio, chromatographic selectivity, detection count and intensity values per study. For all analyses, the full feature table was used which enforces the default feature quality filters (signal-to-noise ratio, or SNR, of 2, and a peak shape > 0.5 defined as goodness of fit to a gaussian model). Orbitrap studies were processed using the autoheight option enabled, while TOF data was processed using a mass accuracy of 25 ppm, a min_peak_height of 1000, a cal_min_peak_height as 3e4, and a min_intensity_threshold of 500. The m/z calibration is performed by aligning all features against a list of resolved mass values from HMDB and PubChemLite and fitting to a linear regression model. Should a systematic shift of m/z values exceed 2 ppm, all m/z values in the dataset are corrected by the regression model. A few datasets produced extra large number of features because MS/MS acquisitions were mixed with MS full scan, which created artificially extra elution peaks. They do not impact the mass registry and are voted down in consensus features. GC-MS and MS/MS data were collected and processed as described previously (Smirnov et al, 2021).

### Data analysis and integration for CSM construction

All features from Orbitrap platforms were used as input to kernel density estimation of m/z distribution. Using a density threshold of 0.05, 48,697 and 40,483 KDE peaks were extracted in positive and negative ion modes, respectively. Each KDE peak produced a consensus m/z value, which became a CSM mass registry. The KDE function was based on Python library statsmodels (version 1.14.0), nonparametric.KDEUnivariate. The KDE peaks were identified using scipy.signal.find_peaks (version 1.13.1). The same KDE parameters were used for TOF data analysis.

Khipu performs pre-annotation by assigning co-eluting features to adduct and isotopologue relations using a generic tree structure based on *a priori* mass delta patterns (Li and Zheng, 2023). The minimum of 5 ppm of m/z value or 0.0005 as absolute value was used for mass delta matching. Detailed parameters are given in the CSM GitHub repository.

CSM features and mass registries are stored as JSON dictionaries. An example of CSM feature is:

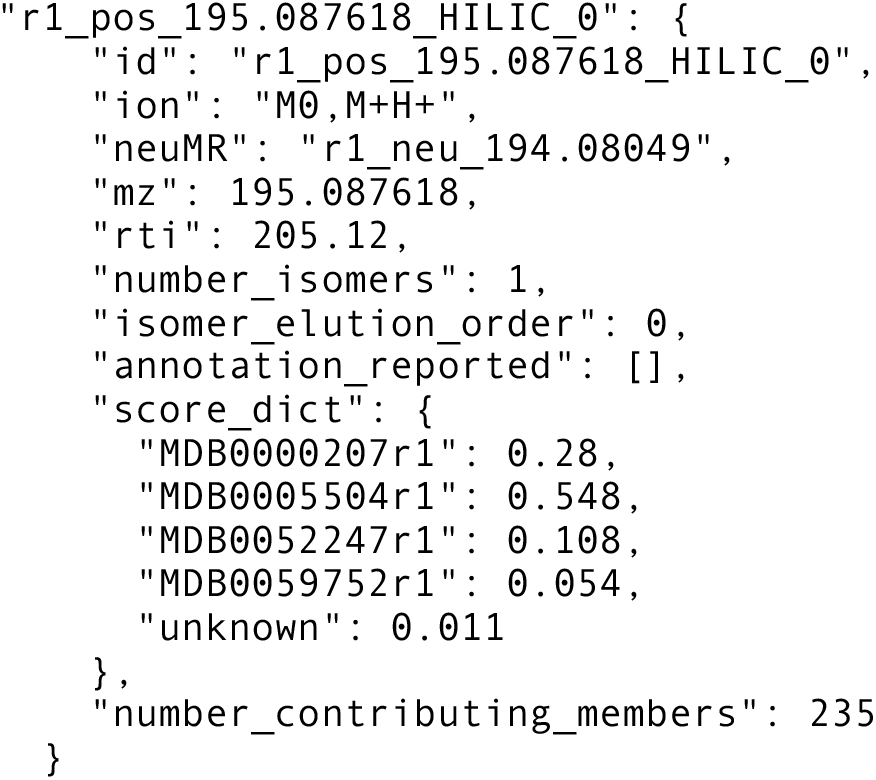

### An example of neutral mass registry is

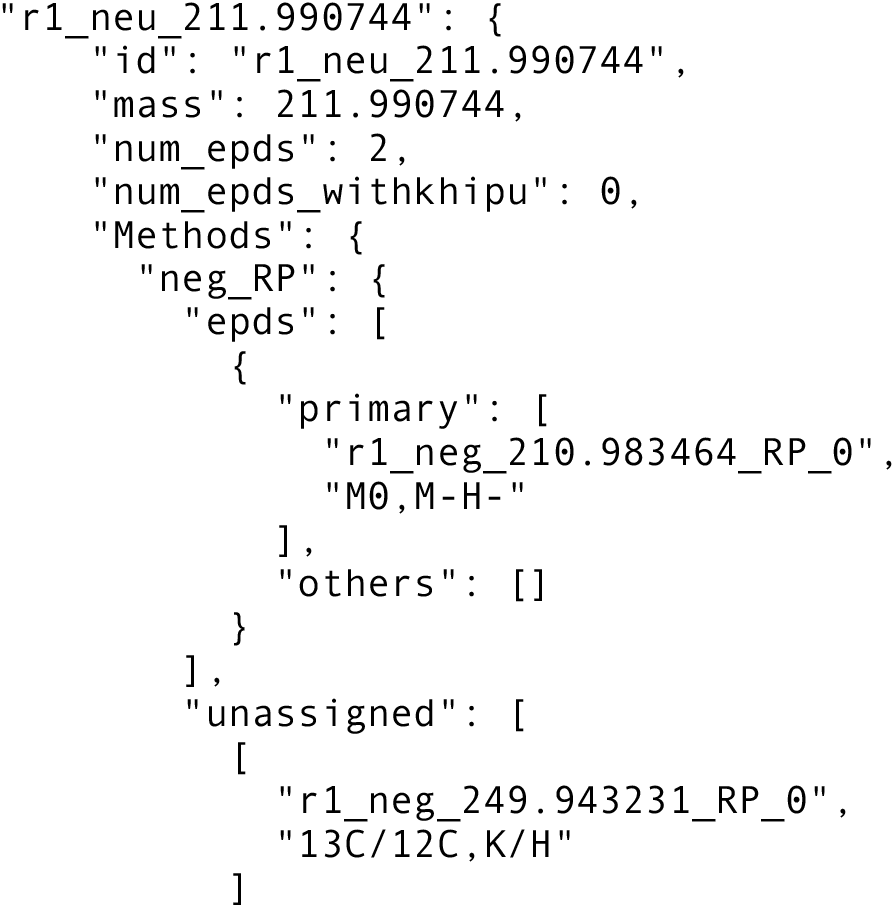
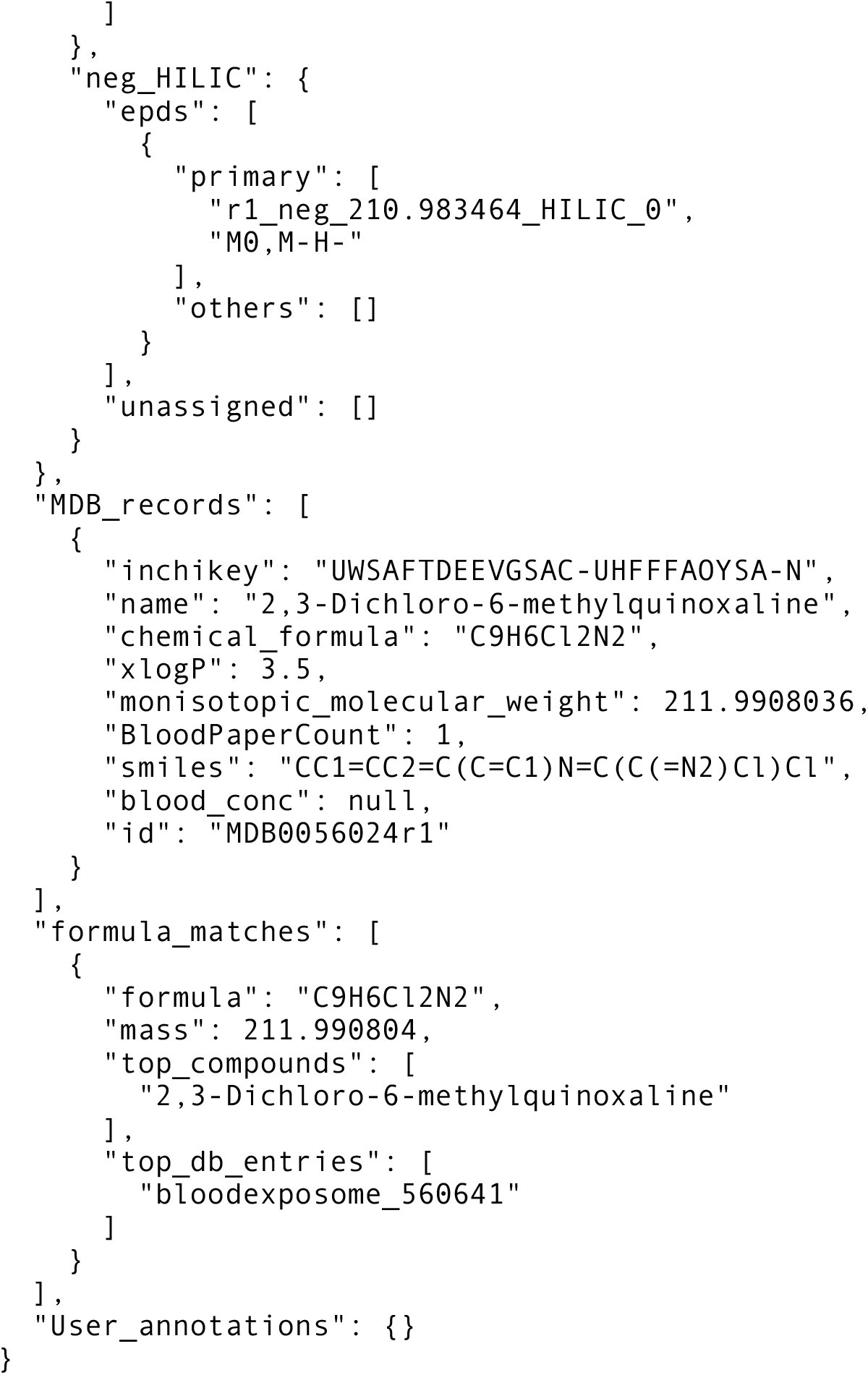

### In-house HZV029 metabolomics data acquisition

Metabolites extraction was carried out by protein precipitation technique using extraction solvent, acetonitrile:methanol (8:1, v/v) containing 0.1% formic acid and isotope labelled Trimethyl-13C3]-caffeine, [13C5]-L-glutamic acid, [15N2]-Uracil, [15N,13C5]-L-methionine, [13C6]-D-glucose and [15N]-L-tyrosine as spike-in controls. 30 μL of plasma sample was taken and 60 μL of extraction solvent was added. Extraction blanks were also prepared to remove features of non-biological origins. All samples were vortexed and incubated with shaking at 1000 rpm for 10 min at 4?C followed by centrifugation at 4?C for 15 min at 20,817 x g. The supernatant was transferred into mass spec vials and 2 μL injected into UHPLC-MS.

All samples were maintained at 4 °C in the autosampler, and analyzed using a Thermo Scientific Orbitrap ID-X Tribid Mass Spectrometer coupled to a Thermo Scientific Transcen LX-2 Duo UHPLC system, with a HESI ionization source, using positive and negative ionizations. The MS settings are: spray voltage, 3500 V; sheath gas, 45 Arb; auxiliary gas, 20 Arb; sweep gas, 1 Arb; ion transfer tube temperature, 325 °C; vaporizer temperature, 325 °C; mass range, 80-1000 Da; maximum injection time, 100 ms. The resolution was set at 120,000 in the HZV029 experiment, 60,000 in the BM21 and SZ22 experiments.

Data were acquired using hydrophilic interaction liquid chromatography (HILIC) positive and reversed phase (RP) negative polarities in full scan mode with mass resolution of 120,000 simultaneously. An AccucoreTM-150-Amide HILIC column (2.6 μm, 2.1 mm x 50 mm) and a Hypersil GOLDTM RP column (3 μm, 2.1 mm x 50 mm) maintained at 45 ºC were used for chromatographic separation. 0.1% formic acid in water and 0.1% formic acid in acetonitrile were used as mobile phase A and B respectively for RP acquisition. 10 mM ammonium acetate in acetonitrile:water (95:5, v/v) with 0.1% acetic acid as mobile phase A and 10 mM ammonium acetate in acetonitrile:water (50:50, v/v) with 0.1% acetic acid as mobile phase B were used for HILIC method. For HILIC acquisition, following gradient was applied at a flow rate of 0.55 ml/min: 0-0.1 min: 0% B, 0.10-5.0 min: 98% B, 5.00-5.50 min: 0% B and 4.5 min for cleaning and equilibration of column. For RP column, following gradient was applied at a flow rate of 0.4 ml/min: 0-0.1 min: 0% B, 0.10-1.9 min: 60% B, 1.9-5.0 min: 98% B, 5.00-5.10 min: 0% B and 4.9 min cleaning and column equilibration. The chromatographic run time was 5 min followed by 5 min washing step after each sample.

### CSM alignment and annotation algorithms

The compound libraries underlying Bar et al (2020) and Pathmasiri et al (2024, ST003110) were verified by m/z and retention time in their corresponding raw data. This also involved studies ‘MTBLS136’, ‘MTBLS204’, ‘MTBLS205’, ‘MTBLS407’ and ‘MTBLS833’, as they share the same library as Bar et al. These compounds established the initial CSM features with confirmed identification; they were then merged with the Blood Compound Database in this project (Suppl Figure 2). InchiKeys were manually matched when other identifiers were unclear.

When a feature table is subjected to CSM annotation, the khipus and primary features are first established within the dataset. The mass tracks of the primary features are then matched to CSM mass registries. If there are multiple features on the mass track or mass registry (isomers), they are aligned by number of consensus features, abundance of the features and elution orders.

All Blood Compound Database entries are considered for annotation, when the mass values agree with a query. The annotation scores are calculated using a framework of factor graph, while not involving statistical learning yet. The CSM features are treated as observed variables and Compound Database entries as domains. An unknown compound and unknown feature are introduced along the presented entities. They are fully connected in a bipartite network with equal weights on the edges. All annotation evidence, including prior identification, known concentration, publication numbers, detection frequency, elution order and partition coefficient, is iterated into a scoring function based on expert heuristics. The scores per compound are normalized and annotation recommendations are made.

### Consensus Retention Time Index

The retention time from the NORMAN database (RP chromatography, Aalizadeh et al, 2021) and RETIP database (HILIC chromatography, Bonini et al, 2020) was introduced into CSM via InchiKey matches to CSM Compound Database. Iteratively, every contributing Orbitrap dataset (from Figure 1) was matched to CSM Compound Database. In every iteration, primary features of clean chromatography (no isomers) were used as anchors to build a regression model for retention time mapping. Finally, the median value of the remapped retention times from all contributing datasets is taken as the consensus retention time index, which was normalized between the range of 0 to 1000.

### Data visualization and web tool

Data visualization is based on the following Python libraries: Matplotlib, Seaborn and lcvk (Li, 2025). The CSM website is hosted on the Google Cloud Platform and is freely accessible at https://metabolomics.cloud/csm. The web application follows a microservice design. The backend processing is Python-based, utilizing Flask to implement an application programming interface (API) to connect the back end to the front end. The front end interactive interface is built using Angular, with elements for interactive data exploration, asynchronously connecting to the back end.

