## Supplementary Figures for "Constructing a consensus serum metabolome"

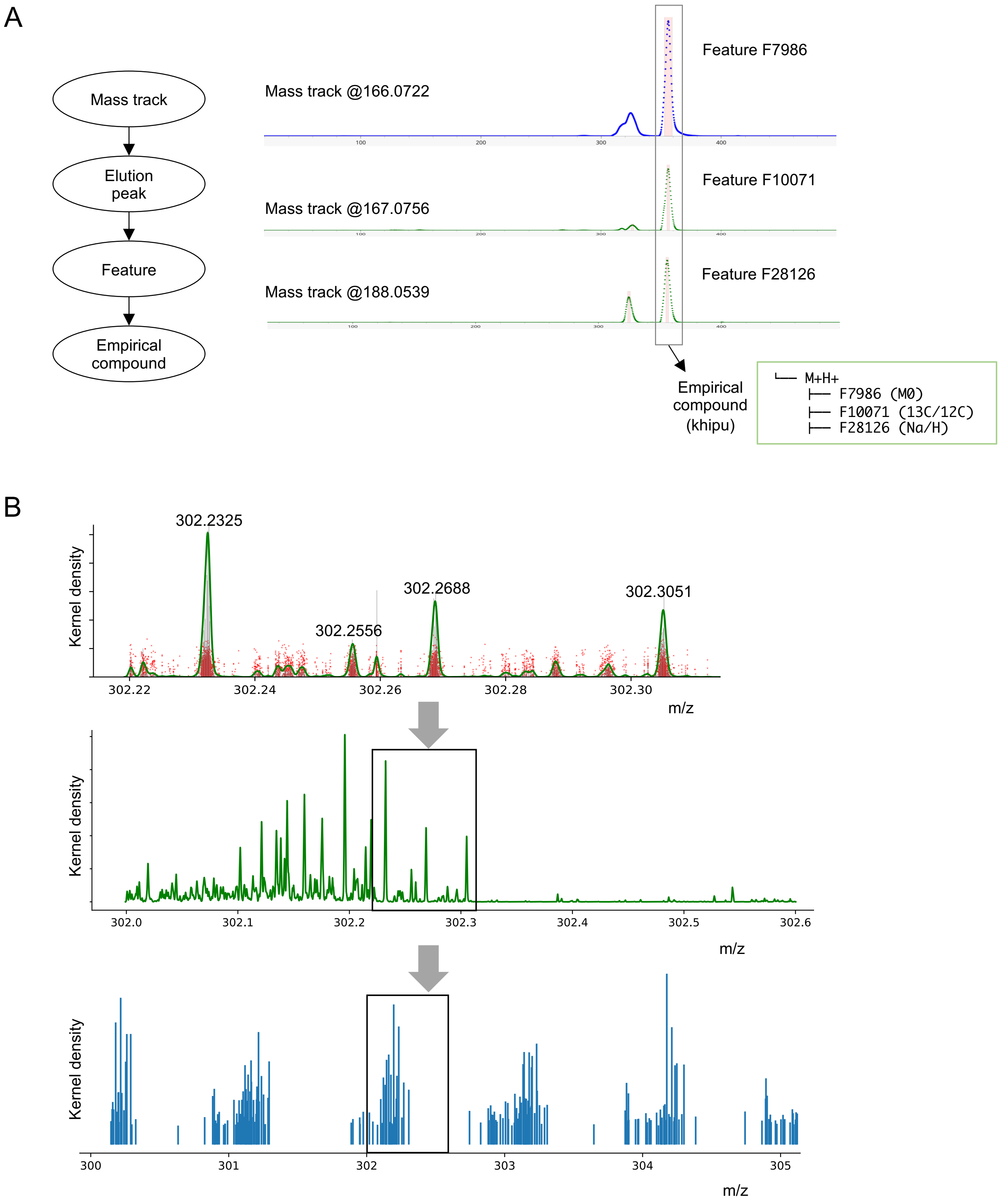


**Supplementary Figure 1: Data models and assembly of consensus mass registries.**

A) A mass track is a unique m/z (mass-to-charge) value measured in an analytical sample (an acquisition file); elution peaks may be detected on a mass track; a feature is defined at the level of a dataset using the same method. A metabolite can be measured by multiple features, including isotopes and adducts, which are grouped into an "empirical compound" by the process of pre-annotation. The implementation of these data structures and related software tools have been described previously (Li et al, 2023; Li and Zheng, 2023; Mitchell et al, 2024a).

B) Peaks of KDE, based on frequency of reported m/z values, are identified as mass registries.

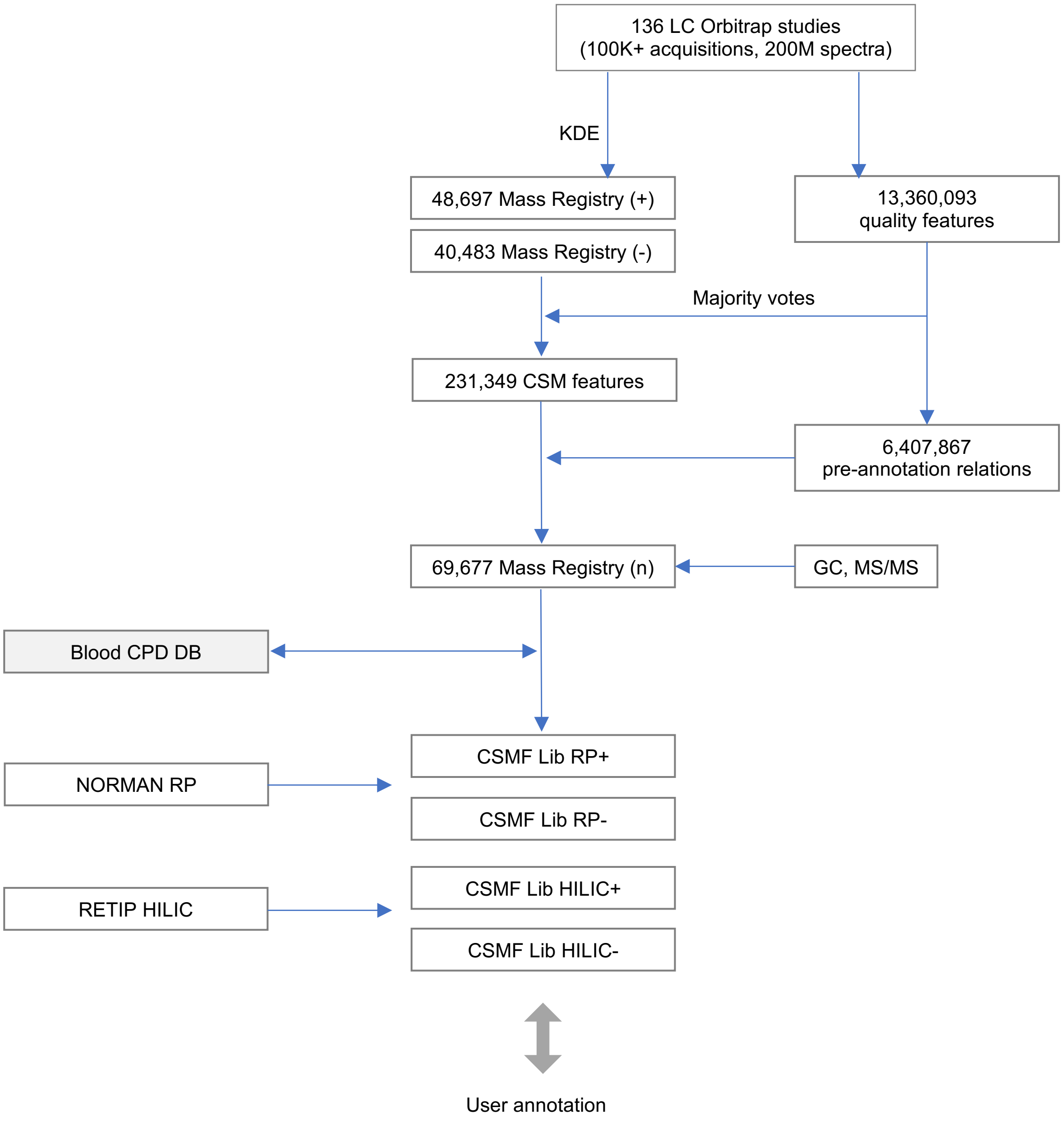


**Supplementary Figure 2: Detailed construction process of CSM.**

CSM features are voted by quality features from individual studies, and linked to mass registries. Feature relationships are determined within each dataset by pre-annotation. Over 6 million ion relationships from all datasets can be transferred into CSM to assign ion types and determine neutral mass values. A neutral mass registry contains consensus features by different methods and their linked annotations. The annotations include curated compound libraries and a generic Blood Compound DB. Retention time indices are computed with integration from compound libraries, NORMAN DB and RETIP library. User data are matched against a method specific library.

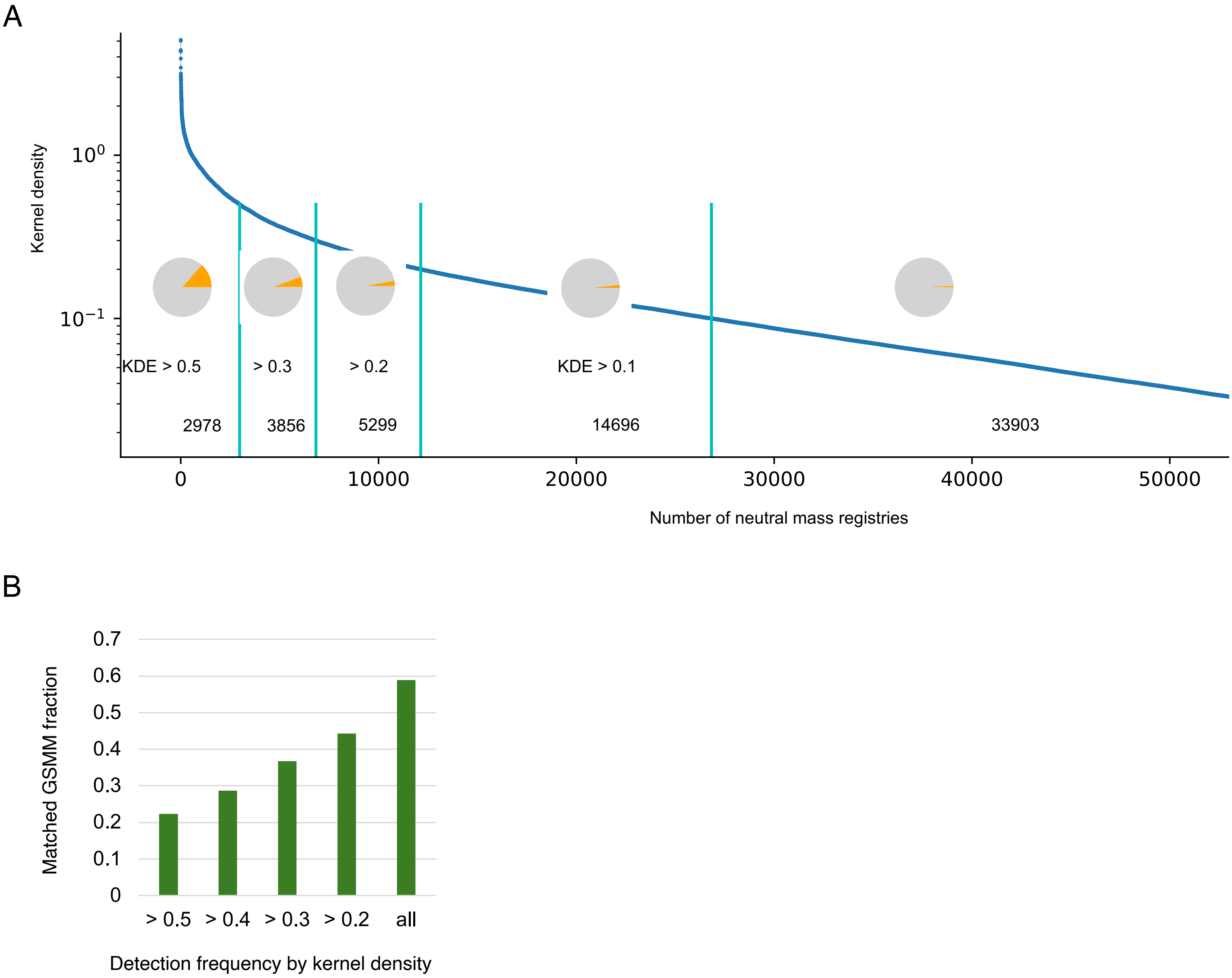


**Supplementary Figure 3: Overlap between CSM and a genome scale metabolic model.**

A) Proportion of CSM features found in the GSSM (Robinson et al, 2020) is dependent on detection frequency.

B) Proportion of GSSM matched to CSM, by detection frequency.

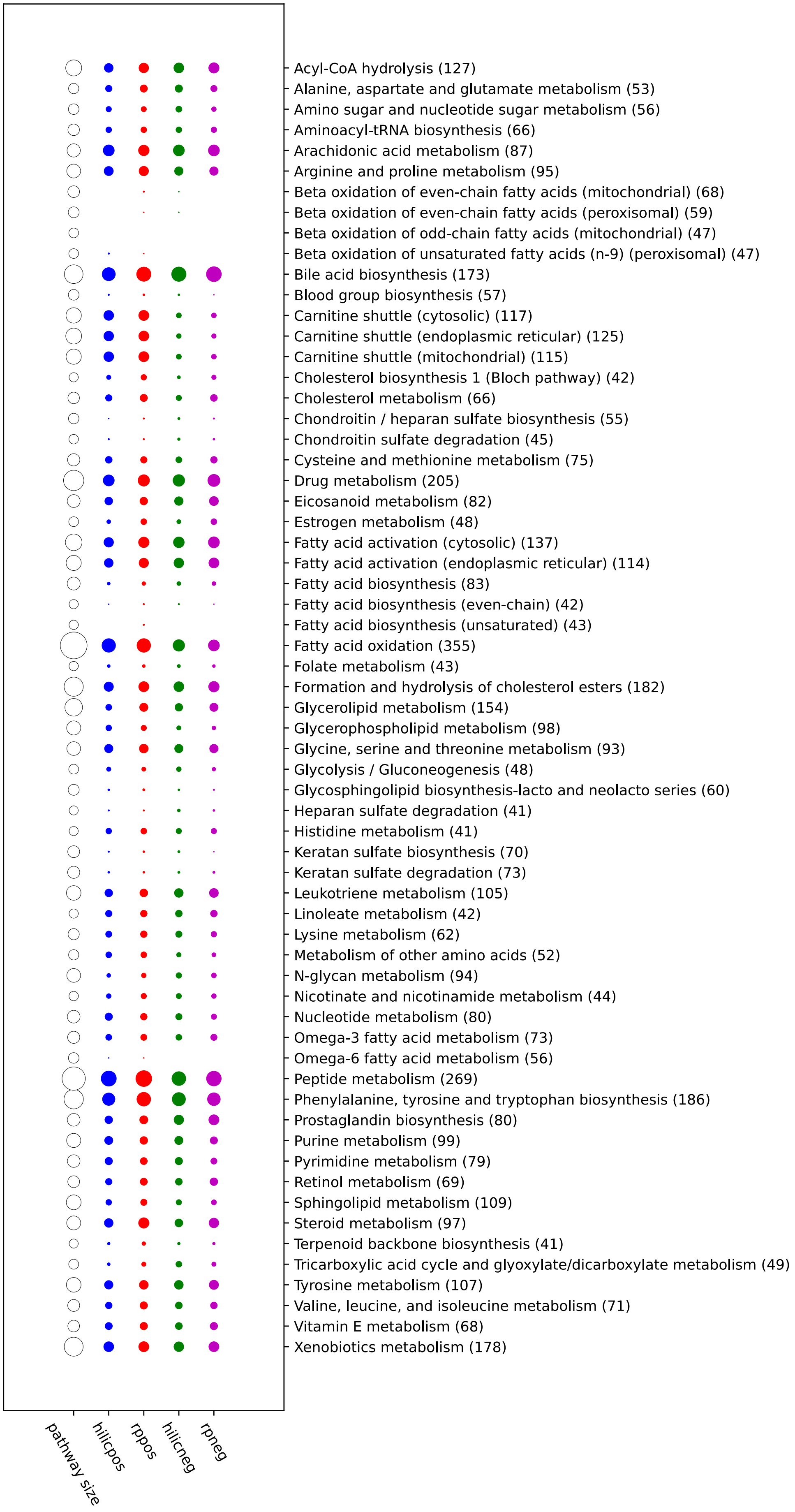


**Supplementary Figure 4. CSM coverage of pathways in human genome scale metabolic model by methods.** Only the top 40 largest pathways are shown here for brevity.

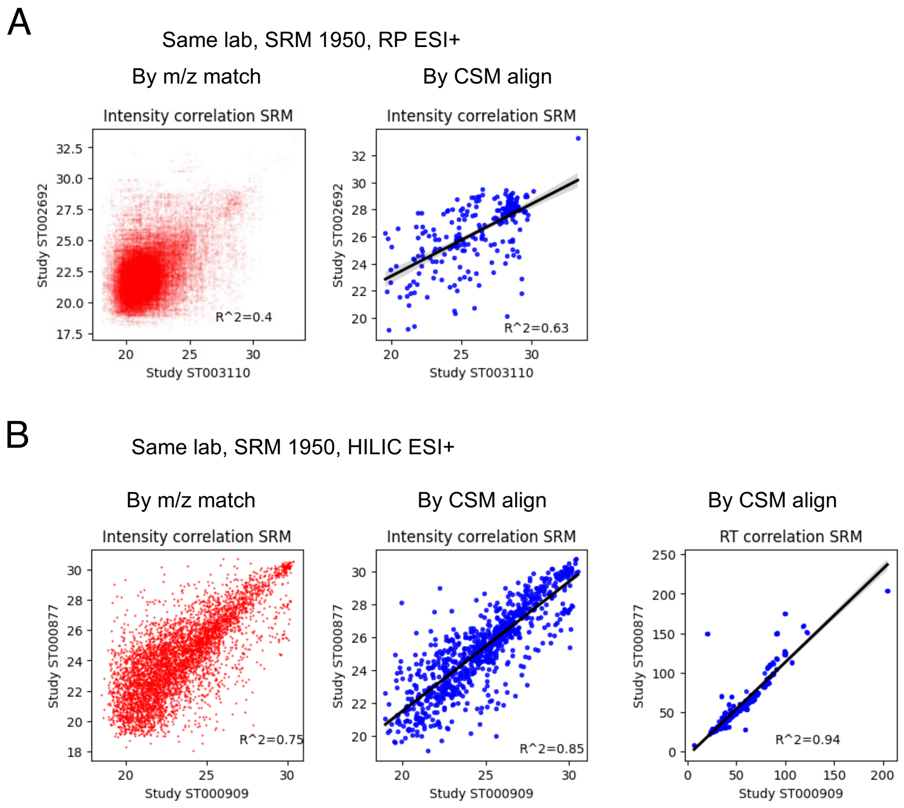


**Supplementary Figure 5: Alignment of studies from the same labs.** Results in red are from features matched by m/z only; blue by CMS alignment.

A) Intensity correlation for the studies in Figure 4C.

B) Two different studies by the same lab using HILIC ESI+ method.

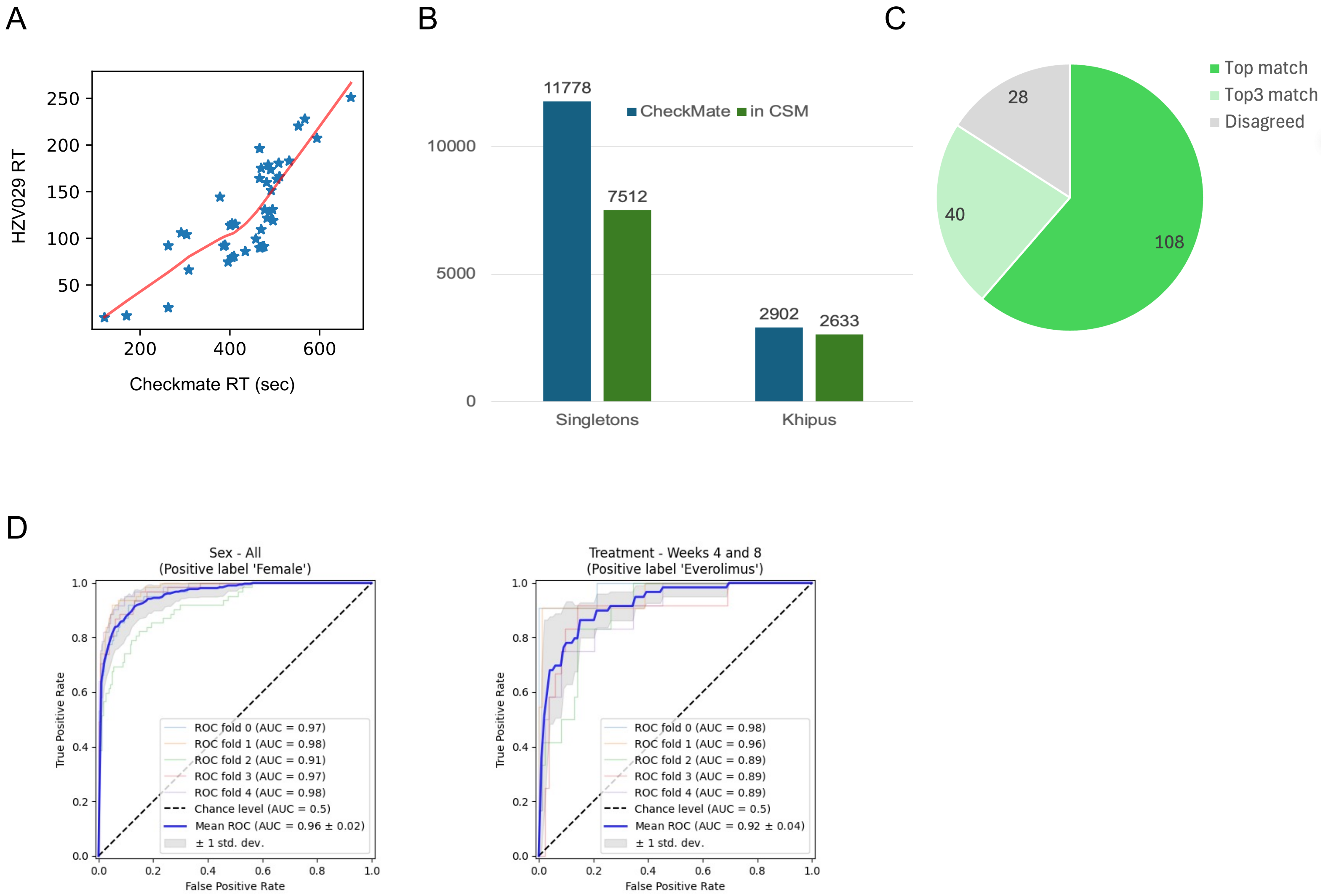


**Supplementary Figure 6: Reprocess, annotation and analysis of CheckMate data.**

A) RT correlation for common confirmed metabolites in both HZV029 and CheckMate (Li et al, 2019).

B) Alignment of features and khipus to CSM.

C) Agreement between CSM named compounds and the authentic compound library from CheckMate.

D) Two different drugs were administered in the study (n = 349/394). SVM models were trained by 5-fold cross validations; prediction results were quantified by Receiver operating characteristic (ROC) curves. Prediction of patient sex is used as control (right).

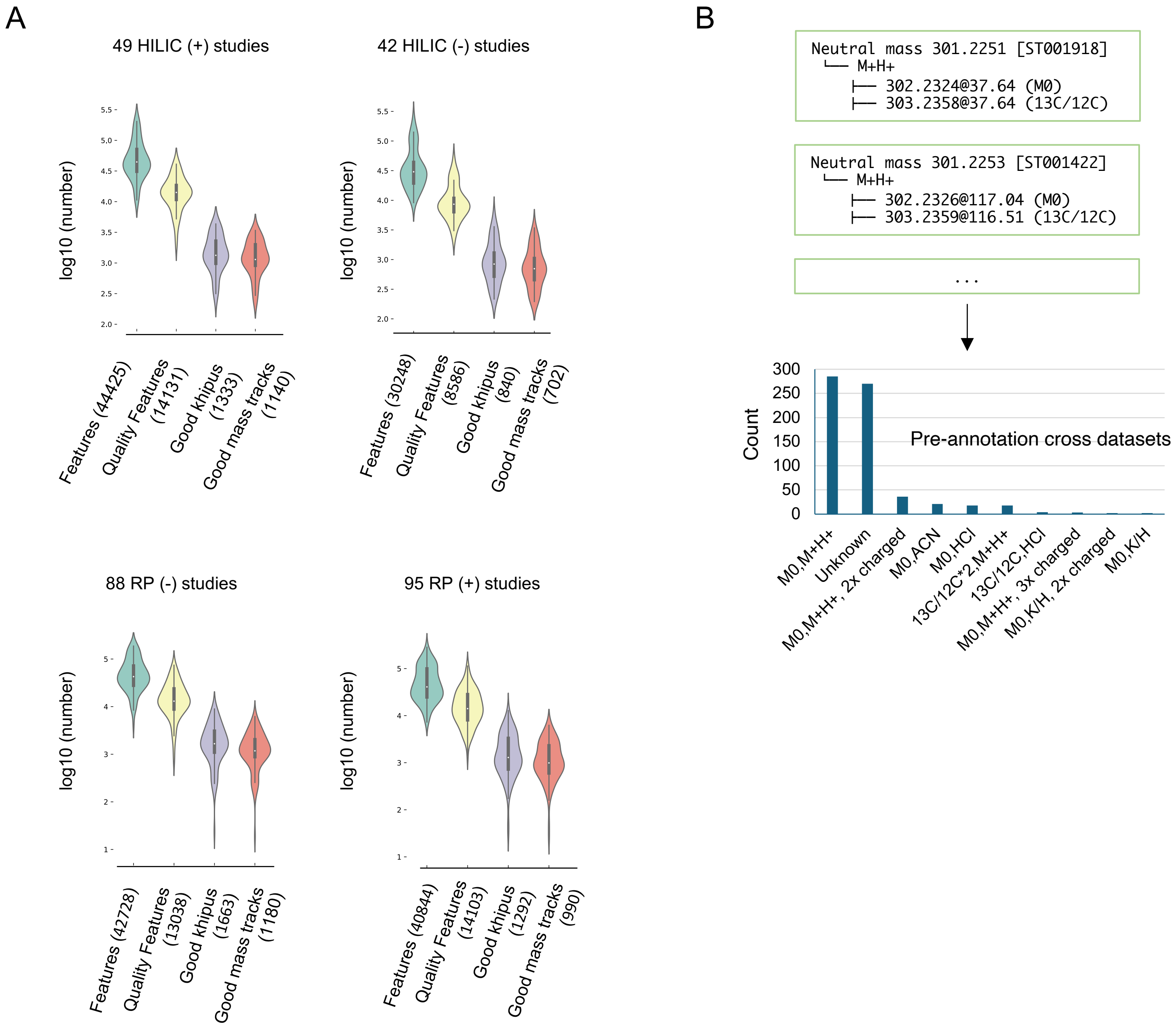


**Supplementary Figure 7: Details of pre-annotation.**

A) Results of all studies by methods. Good features are defined as having SNR > 5 and peak shape > 0.9 when fitting to a gaussian curve. Good khipus have matched 13C/12C pairs. Good mass tracks contain all primary ions of good khipus, thus their ratio indicates prevalence of isomers.

B) Pre-annotations from individual datasets are tallied for a consensus CSM feature.
